# A carbon nanotube optical reporter maps endolysosomal lipid flux

**DOI:** 10.1101/134999

**Authors:** Prakrit V. Jena, Daniel Roxbury, Thomas V. Galassi, Leila Akkari, Christopher P. Horoszko, David B. Iaea, Januka Budhathoki-Uprety, Nina H. Pipalia, Abigail S. Haka, Jackson D. Harvey, Jeetain Mittal, Frederick R. Maxfield, Johanna A. Joyce, Daniel A. Heller

**Author notes:** Contributed equally to this work (Prakrit V. Jena, Daniel Roxbury, Thomas V. Galassi).

## Abstract

Lipid accumulation within the lumen of endolysosomal vesicles is observed in various pathologies including atherosclerosis, liver disease, neurological disorders, lysosomal storage disorders, and cancer. Current methods cannot measure lipid flux specifically within the lysosomal lumen of live cells. We developed an optical reporter, composed of a photoluminescent carbon nanotube of a single chirality, which responds to lipid accumulation via modulation of the nanotube’s optical bandgap. The engineered nanomaterial, composed of short-single stranded DNA and a single nanotube chirality, localizes exclusively to the lumen of endolysosomal organelles without adversely affecting cell viability or proliferation, or organelle morphology, integrity, or function. The emission wavelength of the reporter can be spatially resolved from within the endolysosomal lumen to generate quantitative maps of lipid content in live cells. Endolysosomal lipid accumulation in cell lines, an example of drug-induced phospholipidosis (DIPL), was observed for multiple drugs in macrophages, and measurements of patient-derived Niemann-Pick type C fibroblasts identified lipid accumulation and phenotypic reversal of this lysosomal storage disease. Single-cell measurements using the reporter discerned sub-cellular differences in equilibrium lipid content, illuminating significant intracellular heterogeneity among endolysosomal organelles of differentiating bone marrow-derived monocytes. Single-cell kinetics of lipoprotein-derived cholesterol accumulation within macrophages revealed rates that differed among cells by an order of magnitude. This carbon nanotube optical reporter of endolysosomal lipid content in live cells confers new capabilities for drug development processes and the investigation of lipid-linked diseases.

---

Endosomes and lysosomes are vacuolar organelles responsible for the breakdown of lipids, proteins, sugars, and other cellular materials^1^. The failure to catabolize or export lysosomal contents can result in lysosomal storage disorders (LSDs), a family of approximately 50 diseases characterized by the accumulation of undigested substrates, such as lipids and glycoproteins, within the endolysosomal lumen due to an inherited defect in a single protein^2-3^. Lipid accumulation in the endolysosomal lumen is observed in many LSDs as well as during atherosclerotic foam cell formation^4^, the transition from steatosis to non-alcoholic steatohepatitis^5^, in multiple neurological disorders^6^, cancer^7^, and drug-induced phospholipidosis (DIPL)^8^. The search for small molecule therapeutics against LSDs, as well as our understanding of the aforementioned diseases is hampered by the limited number of tools available to assay lipid content exclusively within the endolysosomal organelles of live cells^9^.

Although multiple classes of sensors and imaging modalities exist to study lipids, current probes are limited in their capabilities. Stains such as LipidTox can detect the general accumulation of lipids within cells^10^ but are not organelle-specific. Fluorophore-conjugated and intrinsically fluorescent lipid analogs are used for analyzing lipid trafficking^10^. Lipid analogs, synthesized by conjugation of a fluorophore to a modified lipid, allow the tracking of uptake and incorporation of lipids into the cell membrane^11-12^. Lipid dynamics can be tracked in live cells using fluorescent proteins fused with lipid-binding domains, however the expressed domains can hamper the native function of the lipid^13^. Environmentally-sensitive fluorophores were recently developed, which can respond to lipid order^14^ in cell membranes undergoing processes such as endosomal maturation^15^, while another family of polarity-sensitive probes, can integrate into lysosomal membranes and detect changes in the overall polarity^16^. The bulk of these technologies are useful for studying lipids present within a biological membrane, but, to the best of our knowledge, no existing probe specifically localizes to the lumen of endolysosomal organelles and reports on the lipid content of its immediate environment.

To develop a biocompatible fluorescent reporter for lipids in the endolysosomal lumen, we investigated the unique physicochemical properties of single-walled carbon nanotubes (SWCNTs). Semiconducting SWCNTs emit highly photostable and narrow-bandwidth near-infrared photoluminescence^17^, which is sensitive to local perturbations^18^. The availability of multiple species with different emission properties can facilitate multiplexed imaging^19^. The SWCNT emission energy responds to the solvent environment^20^ via solvatochromic energy shifts^21^. This response has been used to detect conformational polymorphism^22^ of DNA and the nuclear environment in live cells^23^, as well as microRNA^24^, via shifts down to ≤1 nm. While the self-assembly of lipid derivatives on carbon nanotubes was observed over 14 years ago^25^, the optical response of fluorescent carbon nanotubes to fatty acids has been noted more recently^26^.

Due to their unique applications in biological sensing and imaging applications^27^, the biocompatibility of carbon nanotubes has been a subject of much investigation^28-29^. A recent comprehensive review concluded that the biocompatibility of single-walled carbon nanotubes is dependent on how the nanomaterial sample is processed and functionalized^30^. In particular, multi-walled carbon nanotubes and long single-walled carbon nanotubes or nanotube preparations containing impurities have documented toxic effects on live cells^31^.

Here, we present a biocompatible carbon nanotube optical reporter of lipids within the endolysosomal lumen of live cells. Composed of a non-covalent complex consisting of an amphiphilic polymer and a single (n,m) species (chirality) of carbon nanotube, the reporter exhibits a solvatochromic shift of over 13 nm in response to biological lipids. In mammalian cells, the reporter remains within the endolysosomal pathway and localizes specifically to the lumen of endolysosomal organelles without adversely affecting organelle morphology, integrity, capacity to digest substrates, or cell viability or proliferation. Using near-infrared hyperspectral microscopy, we spatially resolved the solvatochromic response of the reporter to lipids in the endolysosomal lumen and obtained quantitative lipid maps of live cells with sub-cellular resolution. Emission from the reporter identified the lysosomal storage disease Niemann-Pick type C (NPC) in fibroblasts from an NPC patient. Furthermore, the reporter benchmarked treatment, exhibiting a distinct signal reversal upon administration of hydroxypropyl-β-cyclodextrin, a drug that reverses the disease phenotype. Additionally, endolysosomal lipid accumulation was detected using spectroscopy alone, in a 96-well plate format compatible with high-throughput drug screening. Using the reporter, single-cell kinetic measurements in a macrophage model system for lysosomal lipid accumulation identified a sub-population of cells that was both significantly lipid deficient and slower to accumulate cholesterol in the lysosomal lumen. In the context of primary monocyte differentiation into macrophages, we discovered that, as the lipid content in the lumen increases, individual endolysosomal organelles within single cells accumulate lipids at different rates. Thus, our optical reporter enables quantitative imaging and high-throughput measurement of the lipid content in the endolysosomal lumen of live cells.

## Results and Discussion

### Carbon nanotube optical response to lipids

To develop a reporter of endolysosomal lipid content, we first identified a structurally-defined DNA-carbon nanotube complex that responds optically to lipids. Carbon nanotubes were non-covalently functionalized with specific ssDNA oligonucleotides to facilitate separation using ion-exchange chromatography^32^, resulting in suspensions of single-chirality DNA-nanotube complexes. The introduction of low density lipoprotein (LDL), a biochemical assembly composed of lipids and proteins^33^, induced a decrease in emission wavelength (blue shift) that ranged from 0 to 13 nm (Figure S1). The largest solvatochromic response was observed from the (8,6) nanotube spectral band complexed with ss(GT)_6_, a short oligonucleotide which facilitates separation of the (8,6) species (Figure S2)^32^. The isolated ss(GT)_6_-(8,6) complex exhibited absorption bands at 730 nm and 1200 nm and a single photoluminescence emission peak at 1200 nm (Figure 1a, S2-3). Previous work by our lab indicates that the mean length of ss(GT)_6_-(8,6) complexes following the preparation method used is 88.9 nm^34^. We also recently experimentally identified ss(GT)_6_-(8,6) as a sequence-chirality pair that optimally maximized both the high dynamic range of optical modulation by a lipid-like surfactant (SDC) and the stability of the DNA-nanotube complex^35^.

**Figure 1.**
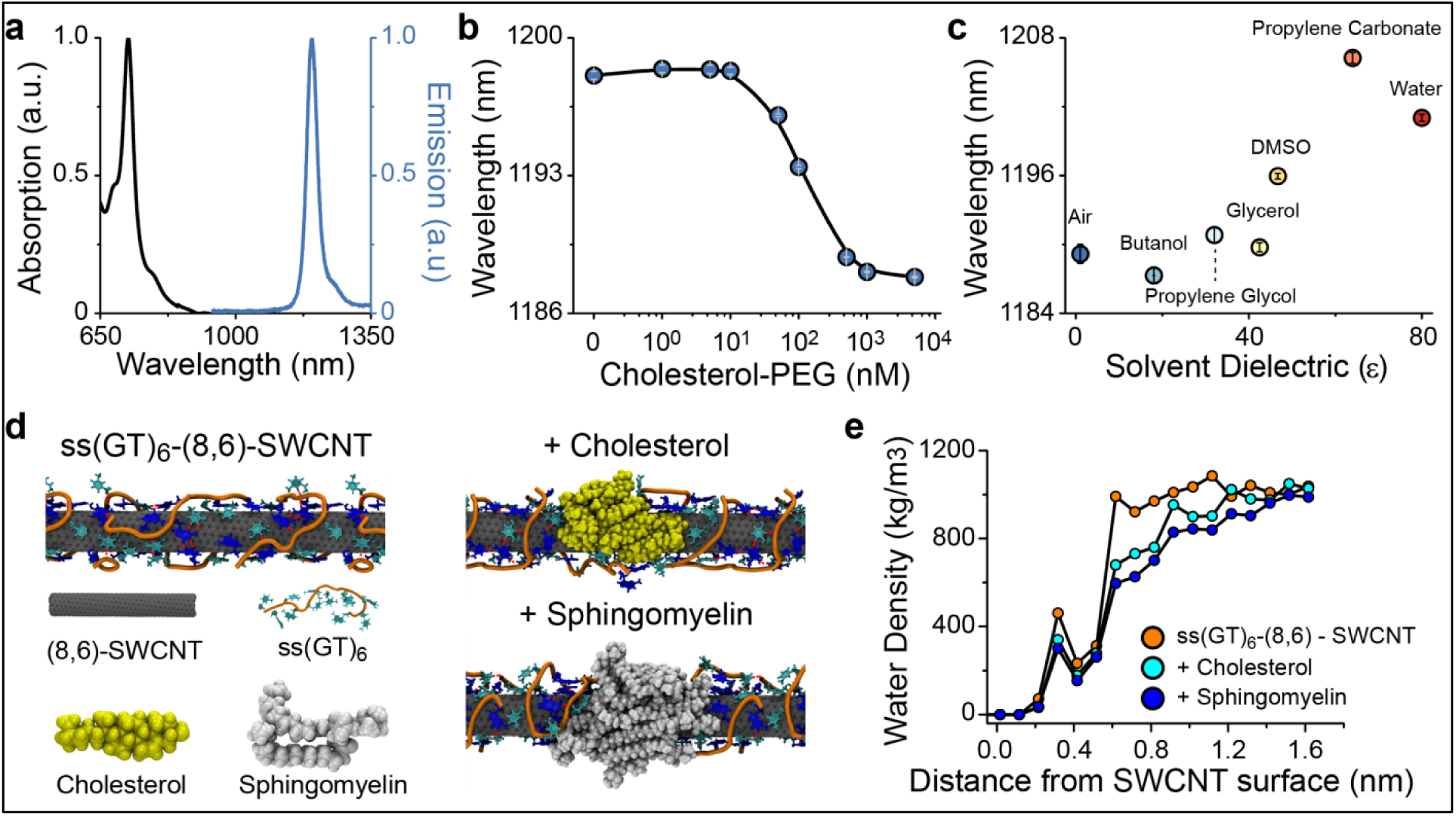
Optical response of carbon nanotube complexes to lipid environments. (a) Normalized absorption and emission spectra of ss(GT)_6_-carbon nanotube complexes purified to isolate the (8,6) species. **(b)** Emission peak wavelength of ss(GT)_6_-(8,6) nanotube complexes in solution as a function of cholesterol-PEG concentration. Error bars are standard error of the mean, obtained from 3 technical replicates performed for each concentration. **(c)** Mean emission wavelength of ss(GT)_6_-(8,6) nanotube complexes exposed to different solvents. Error bars are standard errors of the mean, obtained from 5 technical replicates for each solvent. **(d)** Frames from all-atom molecular dynamics simulations of equilibrated structures of the ss(GT)_6_-(8,6) nanotube complex in water, and the same complex equilibrated in the presence of cholesterol or sphingomyelin. **(e)** Water molecule density as a function of distance from the center of the equilibrated ss(GT)_6_-(8,6) nanotube complex and the same complex equilibrated in the presence of cholesterol or sphingomyelin.

The ss(GT)_6_-(8,6) complex was characterized by measuring the optical response to several classes of biomolecules and water-soluble lipid analogs. Cholesterol-conjugated polyethylene glycol (PEG), a water-soluble analog of cholesterol, induced a ∼ 10 nm decrease in the emission wavelength of the complex at both 2 and 24 hours, while saturating concentrations of BSA, dsDNA from salmon testes, or carboxymethyl cellulose had no measurable effect (Figure S4). The nanotube emission responded rapidly to cholesterol-PEG at 2 hours, but at equilibrium, two different classes of water-soluble lipid analogs elicited equivalent, large blue shifting responses (Figure S5). The optical response of the ss(GT)_6_-(8,6) complex was monotonic and linear over two orders of magnitude of cholesterol concentrations (Figure 1b). A similar response was observed for both ceramide (Figure S6) and low-density lipoprotein (LDL) (Figure S6), indicating the general sensitivity of ss(GT)_6_-(8,6) to both water-soluble lipid analogs and native lipids.

To probe the underlying mechanism governing the response of the ss(GT)_6_-(8,6) complex, we examined the dependence of emission on the dielectric environment^20^. Using near-infrared hyperspectral microscopy^36^, we obtained the emission spectrum from individual surface-adsorbed ss(GT)_6_-(8,6) complexes in 7 different solvent environments (Table S1). The peak emission wavelength of the complexes ranged over 20 nm and exhibited a direct correlation with solvent dielectric constant (Figure 1c) with a Spearman correlation of 0.89, p<0.01. This result suggests that the ss(GT)_6_-(8,6) complex exhibits a distinct solvatochromic response.

For further understanding how lipids interact with the surface of ss(GT)_6_-(8,6) nanotube complexes to induce a solvatochromic shift, we conducted all-atom replica exchange molecular dynamics simulations^37-38^. First, ss(GT)_6_ oligonucleotides were equilibrated on the (8,6) nanotube (Figure S7) to obtain an equilibrium configuration that exhibited a tight association between the ssDNA and nanotube (Figure 1d). Cholesterol molecules were then added and equilibrium was reached after 100 ns (Figure S7). In the resulting configuration, cholesterol bound to exposed regions on the nanotube and induced rearrangement of DNA on the nanotube surface (Figure S8). The combined effect was an 18.7% decrease in the density of water molecules within 1.2 nm of the nanotube surface (Figure 1e). These simulations were repeated with sphingomyelin molecules and a similar reduction in water density was observed (Figure 1d-e). The simulations suggest that lipid binding to the ss(GT)_6_-(8,6) complex reduces the water density on the nanotube surface, thereby lowering the effective local solvent dielectric. As experimentally observed, the lower dielectric environment corresponds to a blue shift of the nanotube emission wavelength (Figure 1c).

We further characterized properties of the ss(GT)_6_-(8,6) optical response to cholesterol. The emission shift on cholesterol addition to surface-adsorbed complexes was rapid (under 2 minutes, limited by the hyperspectral instrument acquisition time, Figure S9). Sodium deoxycholate, a surfactant and water-soluble cholesterol analog, was added and subsequently removed from the surface-adsorbed complexes, demonstrating that the wavelength shift on analyte binding is intrinsically reversible (Figure S9). Furthermore, in an acidic environment, the response of the nanotube complex to lipids was similar to that at a neutral pH (Figure S9).

Overall, the characteristics of the ss(GT)_6_-(8,6) complex suggest that it can function as a reporter of endolysosomal lipid accumulation in live cells. When prepared via previously described methods^34^ suspensions of ss(GT)_6_-(8,6) consist of short (∼90 nm), singly dispersed nanotubes that are relatively free of impurities, and non-covalently functionalized with biocompatible single stranded DNA. This minimizes the key parameters of SWCNT cellular toxicity^30^, a topic that is assessed below. The sample length distribution lies between ultra-short (50 nm) and short (150 nm) nanotubes, which maximizes cellular uptake of fluorescent nanotubes while minimizing bundling within cells^39^. The observed brightness of structurally-sorted ss(GT)_6_-(8,6) is intrinsically higher than unsorted DNA-nanotube sensors, as on-resonance excitation at 730 nm efficiently excites every nanotube present. Additionally, this particular sequence-chirality pair is relatively stable, and retains its structural integrity under surfactant exchange^35^. The structure, stability, and brightness of ss(GT)_6_-(8,6), combined with its sensitivity and specificity to lipids over other classes of biomolecules, suggests that it may be applied to live-cell measurements of lipid accumulation.

### Uptake and localization of DNA-SWCNTs to the endolysosomal lumen

Although past work suggests that DNA-SWCNTs incubated with cells are taken up via energy dependent processes and localize to the endolysosomal lumen, this has not been assessed quantitatively or on a single-organelle level^40-42^. We quantitatively assessed the interaction of ss(GT)_6_-(8,6) complexes with mammalian cells using near-IR and visible fluorescence microscopy. Macrophages were incubated in complete 10% FBS-supplemented cell culture media with 0.2 µg/mL of ss(GT)_6_-(8,6) complexes at 37 °C for 30 minutes, before washing with fresh, complex-free media. For the conditions used in our experiments, this concentration corresponds to approximately 39 pM of ss(GT)_6_-(8,6)^34^. The cells exhibited bright near-IR emission, indicating that nanotubes were strongly associated with the cells (Figure S10). We also measured the uptake of the ss(GT)_6_-(8,6) complexes as a function of temperature, and quantified the nanotube emission intensity associated with cells. Incubation at 4 °C resulted in 10-fold lower intensity than incubation at 37 °C (Figure S11), indicating that the complexes had been internalized by the cells via an energy dependent process. These results are consistent with previous reports of the energy-dependent uptake of ssDNA-nanotube complexes^43^ via endocytosis^40-41^. The nanotube emission, quantified from over 700 individual cells incubated with the complex at 37 °C, followed a normal distribution, suggesting a relatively homogeneous uptake of the ss(GT)_6_-(8,6) complexes by the cells (Figure S10).

To determine the localization of ss(GT)_6_-(8,6) complexes following uptake, we conducted a series of imaging experiments in macrophages (RAW 264.7 cell line and bone-marrow derived monocytes) and epithelial (U2OS) cells. The ss(GT)_6_ oligonucleotide was covalently labeled with visible fluorophores (Cy3, Cy5, or Alexa-647) to prepare fluorophore-labeled ss(GT)_6_-nanotube complexes. Following internalization of the complexes by macrophages, we acquired emission from the Cy3 dye and nIR emission from the nanotubes in the same imaging field. The colocalization between the signals, observed on two different detectors, indicates that emission from a fluorophore conjugated to the ssDNA on the nanotube is a reliable indicator of nanotube location (Figure S12). Next, we colocalized the Cy5-DNA-SWCNT fluorescent complexes with Lysotracker Green, a fluorescent probe which accumulates in endolysosomal organelles (Figure 2a, S13). A quantitative analysis using an unbiased auto-thresholding approach indicated significant colocalization between the Cy5 and Lysotracker Green emission (Pearson coefficient of 0.92 ± 0.036, Manders split colocalization coefficient of 0.95 ± 0.018), suggesting that the nanotube signal was contained within endolysosomal organelles. Concomitantly, the nIR emission from the nanotubes localized to the same regions of the cell as the visible emission from Lysotracker, further supporting the endolysosomal localization of the nanotubes (Figure S14).

**Figure 2.**
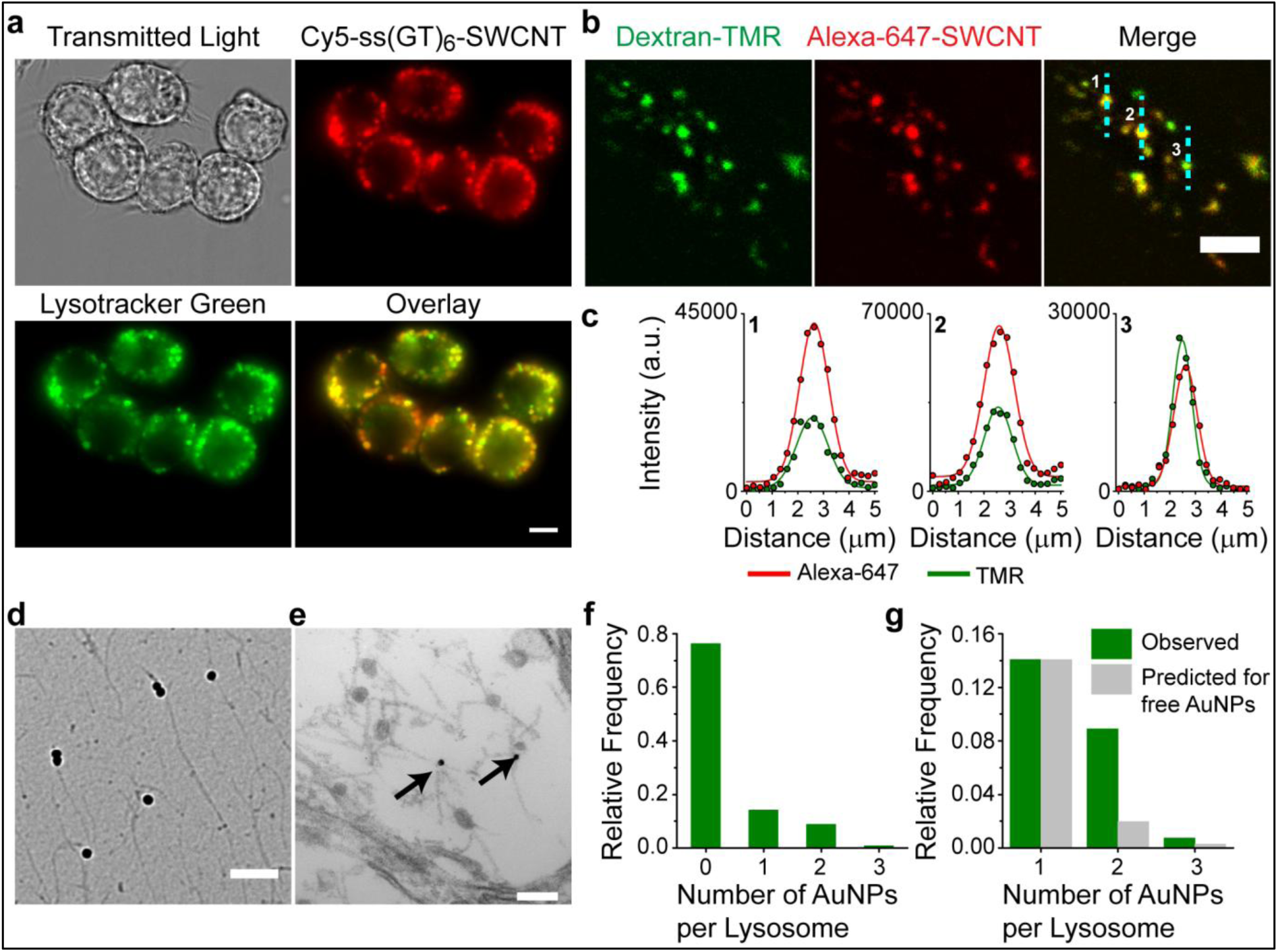
Localization of DNA-SWCNT to endolysosomal organelles. (a) Representative fluorescence microscopy images of Cy5-labeled DNA-SWCNT complexes (red) and LysoTracker (green) in live cells. Scale bar = 10 µm. **(b)** Representative confocal images of TMR-dextran (green) and Alexa-647-SWCNT (red) in U20S-SRA cells. Lines (cyan) denote cross sections from the images extracted for further analysis in (c). Scale bar = 10 µm. **(c)** Intensity profiles of TMR (green) and Alexa 647 (red) fit with Gaussian functions. **(d)** Representative TEM images of AuNP-SWNCT complexes images on the TEM grid. Scale bar is 250 nm. **(e)** Representative TEM image of an AuNP-SWCNT complex within an endolysosomal organelle. Scale bar is 100 nm. **(f)** Relative frequency histogram of AuNP-SWCNT complexes per endolysosomal organelle. **(g)** Relative frequency histograms comparing the experimentally observed and predicted numbers of AuNPs per endolysosomal organelle if AuNPs were not attached to SWCNT complexes.

We next assessed the localization of the nanotubes within the lumen of individual endolysosomal organelles, by pulsing the cells with fluorescent (TMR) 10,000 MW dextran, a polymer which accumulates in the endolysosomal lumen and does not degrade^44^. Following overnight incubation, the cells were maintained in dextran-free media for 3 hours, before Alexa-647 labeled nanotube complexes were introduced to the cell media for 30 minutes and then washed away. One hour later we performed high magnification confocal microscopy in the live cells. An analysis of over 40 cells (Figure 2b, S15) indicates that 50% TMR dextran labeled endolysosomal organelles colocalized with Alexa 647-nanotube complexes, suggesting that within an hour following their addition, the nanotubes had been transported to the dextran-loaded endolysosomal organelles. We extracted line intensity profiles of TMR and Alexa 647 emission and fit them with Gaussian functions (Figure 2c, S15). The single Gaussian intensity distributions of both fluorophores overlapped significantly, with centers that colocalized with diffraction-limited resolution. This result suggests that the nanotubes localized to the same region of endolysosomal organelles as dextran, which is known to remain in the endolysosomal lumen^44^.

To further confirm the presence of DNA-SWCNTs in the endolysosomal organelles, TEM analysis was performed. As single-walled carbon nanotubes, composed of only one layer of cylindrical graphene, do not have sufficient electron density to be visible by TEM in cells, we used gold-labeled DNA-nanotube complexes to perform the first incidence of gold-enhanced TEM imaging of individualized SWCNTs in mammalian cells. Citrate-capped gold nanoparticles (∼ 10 nm diameter) were conjugated to thiolated ssDNA-nanotube complexes^45^. Unbound gold nanoparticles were removed via centrifugation. Images of the gold nanoparticle-nanotube complexes (AuNP-SWCNT) deposited directly onto a TEM grid (Figure 2d, S16a) confirmed that all gold nanoparticles were attached to carbon nanotubes. We then incubated RAW 264.7 macrophages with 1 µg/mL of the gold-labeled nanotubes, and fixed the cells for TEM imaging after removing free gold-nanotube complexes from the solution. In the cells, the gold nanoparticles were clearly visible as dark circles within endolysosomal organelles (Figure 2e, S16b-c). We quantified the number of AuNP-SWCNTs within each endolysosomal organelle (Figure 2f). From the relative frequency distribution, we found the probability that an endolysosomal organelle had one gold nanoparticle was 0.14. If two gold nanoparticles were to independently localize into the same vesicle, we calculated that the probability would be approximately 0.02 (0.14 x 0.14 = 0.02). In contrast, the experimentally determined number of endolysosomal organelles with two AuNP was four times higher (Figure 2g), suggesting that if two AuNP were observed within one endolysosomal organelle, then the two nanoparticles were statistically likely to be linked to each other via a nanotube. This, combined with the removal of free gold nanoparticles via centrifugation and the TEM images showing that all visualized AuNPs were attached to SWCNTs, suggests that the AuNPs within endolysosomal organelles were part of AuNP-SWCNT complexes.

To determine the long-term fate of the complexes, we acquired nIR movies of ss(GT)_6_-(8,6) complexes within macrophages at 6, 24 and 48 hours after the initial 30 minute incubation (Supplementary Videos 1, 2, and 3). At each time point, emission from the complexes exhibited both passive diffusion and directed, linear movements consistent with the active transport of lysosomes along microtubules^46^, suggesting that the nanotube complexes remain within endolysosomal organelles. The emerging view, from our series of experiments, indicates the efficient uptake of DNA-nanotube complexes via endocytosis, rapid transport to the late endosomes and lysosomes, and stable localization to the lumen of endolysosomal organelles.

### Biocompatibility of DNA-SWCNT complexes in mammalian cells

We conducted several experiments to assess the degree to which ss(GT)_6_-(8,6) perturbed cells and endolysosomal organelles in order to determine whether this may be a complication of its use as a live-cell sensor. Using an annexin V and propidium iodide assay, we found that, at its working concentration (0.2 µg/mL), ss(GT)_6_-(8,6) did not affect cell viability or proliferation (Figure S17). To determine if DNA-SWCNT complexes altered the morphology of endolysosomal organelles, AuNP-SWCNT complexes were incubated with RAW 264.7cells for 30 minutes before fixing and preparing for TEM imaging 6 hours later (Figure 3a). Analysis of the size, diameter and aspect ratio of endolysosomal organelles from the TEM images show no statistical differences between control macrophages and macrophages incubated with 1µg/mL complexes (Figure 3b, S18). Endolysosomal organelles in which gold-nanotubes were explicitly detected also displayed similar morphology to controls (Figure 3c, S19). At this elevated concentration of complexes (1 µg/mL), we also did not observe a change in endolysosomal membrane permeabilization (Figure S20).

**Figure 3.**
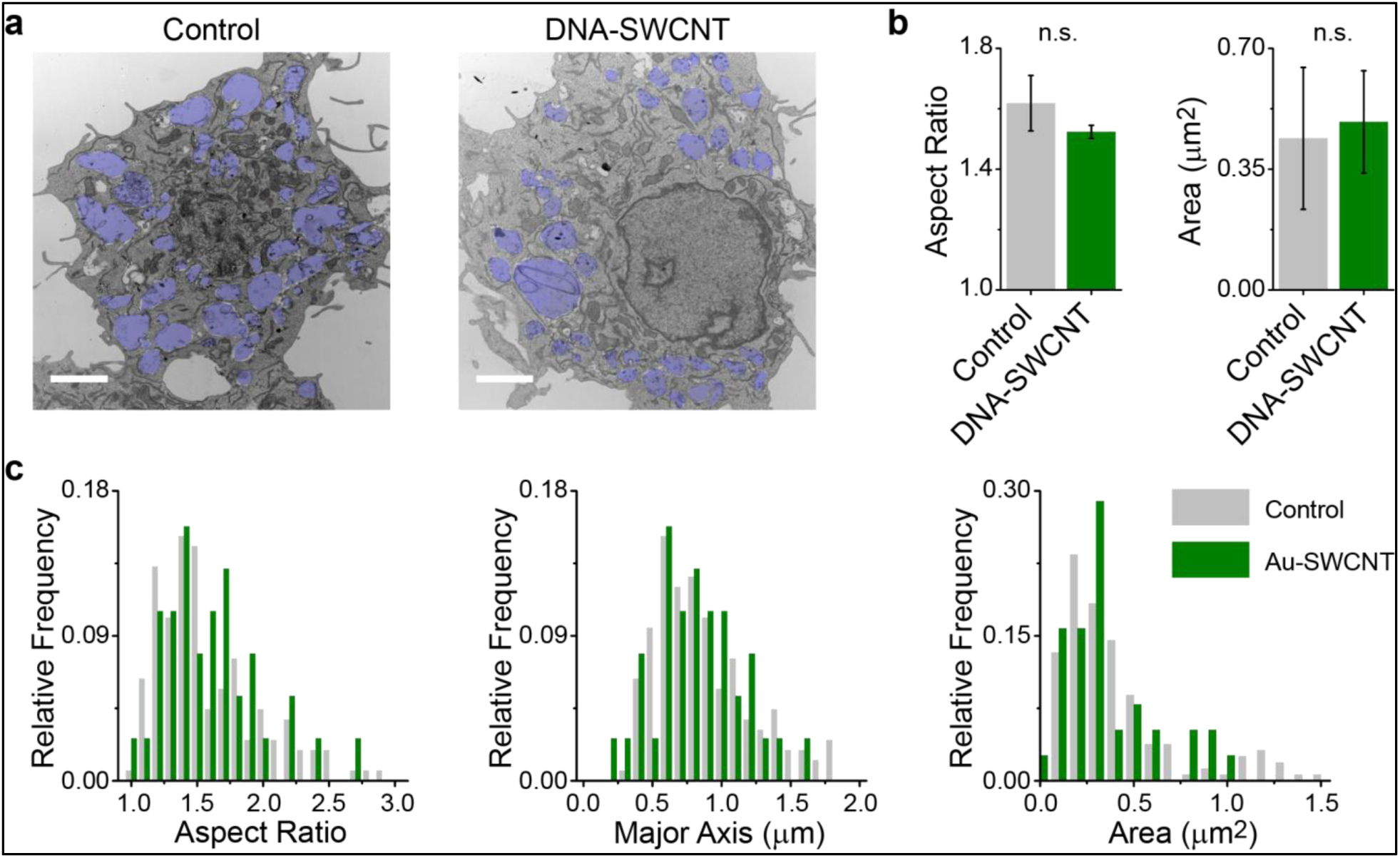
Ultra structural analysis of endolysosomal organelles. (a) Representative TEM images of cells that were untreated or incubated with 1 µg/mL of AuNP-SWCNT complexes. Endolysosomal organelles are shaded blue, scale bars are 2 µm. **(b)** A comparison of the mean aspect ratio (left) and area (right) of endolysosomal organelles. Error bars are standard deviation and mean values were compared with an unpaired t-test. **(c)** Histograms of the distribution of the aspect ratio, major axis, and area of endolysosomal organelles from control cells and endolysosomal organelles containing AuNP-SWCNT complexes.

We next assessed whether DNA-SWCNTs perturbed the ability of endolysosomal organelles to maintain their pH gradient. This was done via a confocal imaging study using LysoTracker, a lysomotropic fluorescent probe that accumulates in acidic vesicles. Human bone osteosarcoma epithelial cells transfected with type A scavenger receptors (U2OS-SRA)^47^ were treated with 1 µg/mL (five times higher than the working concentration) fluorophore-labeled DNA-nanotube complexes (Alexa 647-SWCNT) and fixed and stained with LysoTracker. Using fluorescence confocal microscopy, we imaged both Lysotracker and Alexa 647-SWCNT emission from the endolysosomal organelles (Figure 4a, S21). If DNA-SWCNTs prevented endolysosomal organelles from maintaining a pH gradient, we would expect endolysosomal organelles containing the DNA-SWCNTs to show decreased levels of LysoTracker signal. This was not the case as quantification of LysoTracker fluorescence from organelles both with and without DNA-SWNCT showed no significant difference in LysoTracker intensity (Figure 4b-c, S21).

**Figure 4.**
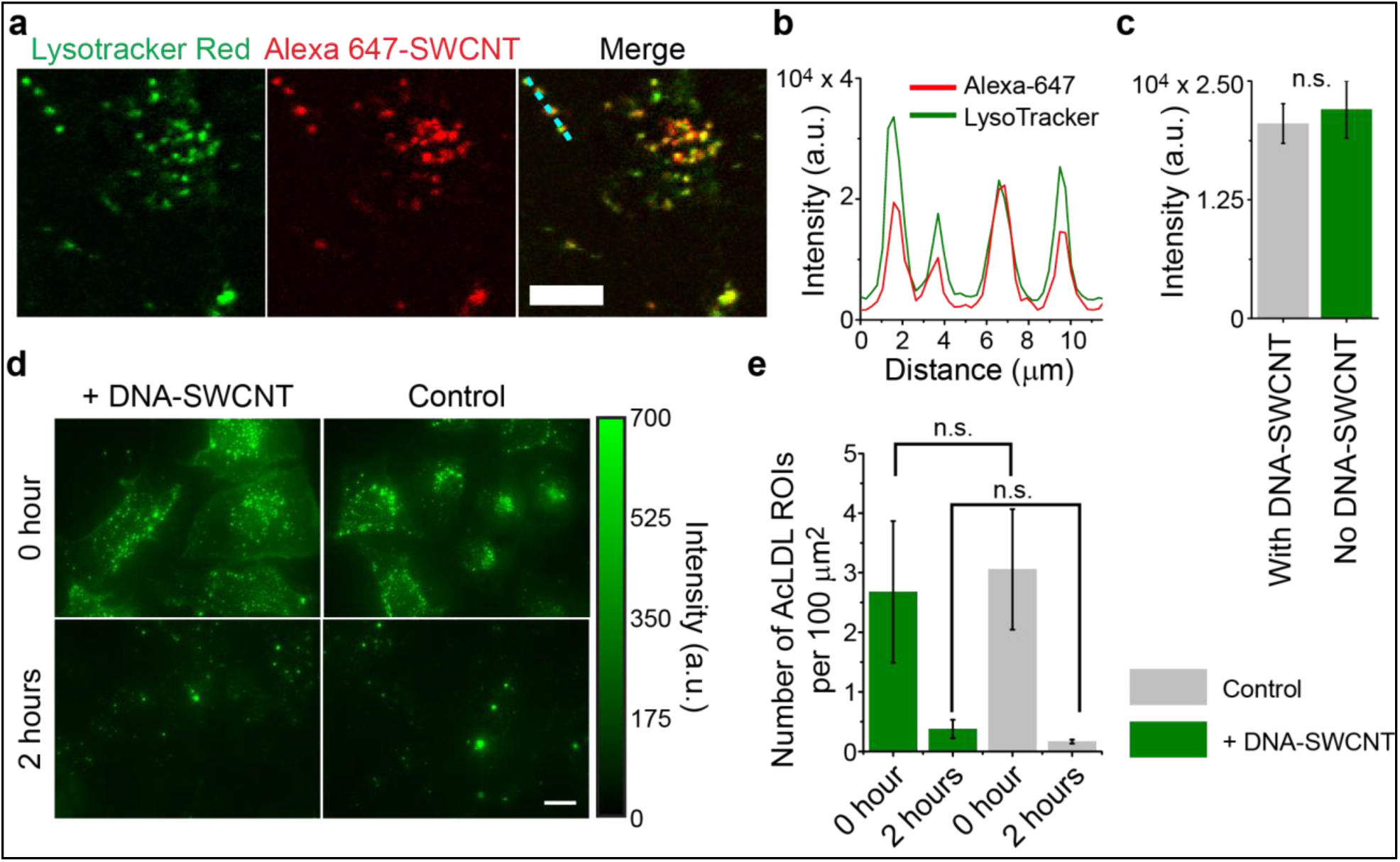
Assessing the effects of DNA-SWCNT on endolysosomal function. (a) Representative confocal images of LysoTracker (green), Alexa 647-SWCNT (red) and a merged image of the two in U2OS-SRA cells Scale bar is 10 µm. **(b)** Intensity profiles of the two fluorophores along the dashed line (cyan) in the merged image. **(c)** Mean intensity of LysoTracker in endolysosomal organelles that contain Alexa 647-SWCNT emission and those that did not. Error bars are standard deviation. Mean intensity was compared with an unpaired t-test (n=9 images per channel). **(d)** Representative epifluorescence images of Alexa 488-AcLDL (green) in U2OS-SRA cells, at 0 and 2 hours after the addition of acetylated LDL, or in control cells. Scale bars are 10 µm. **(e)** Number of AcLDL ROIs per 100 µm^2^. Error bars are standard deviation, obtained from 10 images per condition. Data were compared using a one way ANOVA with Tukey’s multiple comparison test.

To ensure that we could ascertain meaningful results from a DNA-SWCNT based reporter, we also investigated the effect of DNA-SWCNTs on the ability of endolysosomal organelles to hydrolyze lipoprotein molecules. We treated U2OS-SRA cells with 1 µg/mL of Alexa 647-SWCNT for 30 minutes before incubation in fresh media for two hours. To induce the rapid uptake of lipoproteins, we then introduced 50 µg/mL of Alexa 488-labeled acetylated LDL (Alexa 488-AcLDL) to the cell media. Epifluorescence images showed a stark decrease in AcLDL puncta in both control and nanotube-treated cells between zero and two hours of AcLDL addition (Figure 4d, S22). Quantification of AcLDL regions of interest (ROIs), normalized by cell area showed that there was no significant difference in the number of AcLDL ROIs per 100 µm^2^ between cells treated with the nanotubes and the control cells at both 0 and 2 hours (Figure 4e), suggesting that DNA-SWCNTs did not perturb the hydrolysis of lipoproteins. A lipid biochemical assay showed no significant differences in the cholesterol or total lipid content of cell fractions in control and nanotube-containing cells (Figure S23). LipidTox imaging and the expression of LDL receptor (LDLr), also suggested that DNA-SWCNT complexes did not significantly alter lipid processing in cells (Figure S24-25).

The above results suggest that singly-dispersed DNA-SWCNTs that have been separated from large nanotube bundles and carbonaceous impurities via ultra-centrifugation did not adversely affect cell viability or proliferation, or organelle morphology, integrity, or capacity to digest lipoproteins. Moreover, as electronic structure and chirality of a nanotube sample were not found to directly affect toxological impact^48^, our conclusions on biocompatibility likely hold for other relatively short DNA-single walled carbon nanotube complexes composed of different DNA sequences and chiralities.

### Detecting lipid accumulation in the endolysosomal lumen of live cells

We investigated whether the ability of ss(GT)_6_ –(8,6) to detect lipids in vitro could be translated to the endolysosomal organelles of live cells (Figure 5a). RAW 264.7 macrophages were incubated with the complexes for 30 minutes in complete 10% serum-supplemented media at 37° C and washed with fresh media. To induce lysosomal accumulation of free cholesterol, we prepared cells that were pre-treated with U18666A^49^, a compound that inhibits the action of Niemann-Pick C1^50^, a membrane protein that effluxes free cholesterol out of the lysosomes (Figure 5a), to mimic the Niemann-Pick C1 disease phenotype. Cells were also pre-treated with Lalistat 3a2^51^, an inhibitor of lysosomal acid lipase (LAL), which is the central enzyme that hydrolyzes low density lipoproteins (Figure 5a)^52^. Inhibition of LAL leads to the accumulation of esterified cholesterol, which is observed in Wolman’s disease.

**Figure 5.**
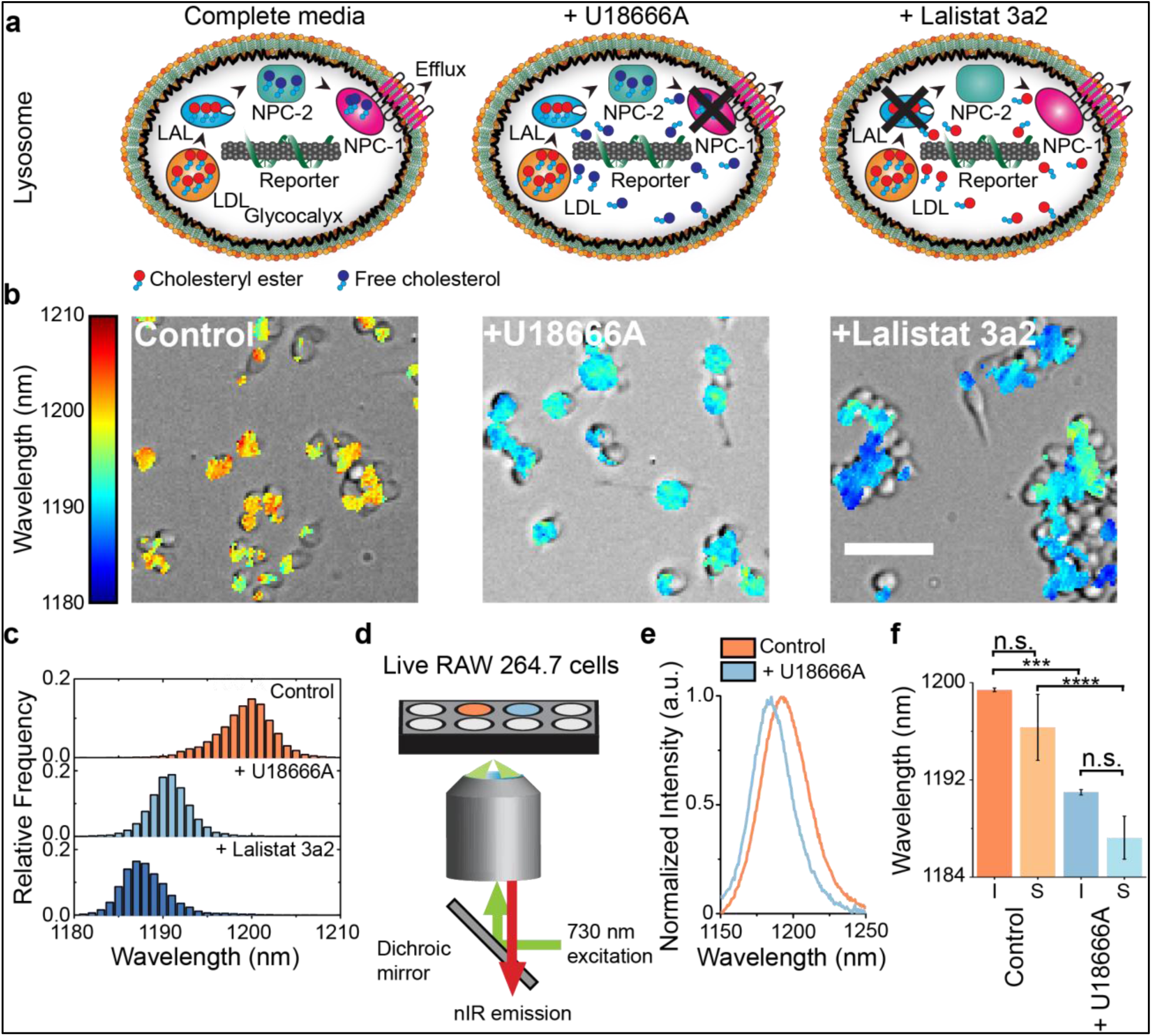
Detection of endolysosomal lipid accumulation in live cells. (a) Schematics of the ss(GT)_6_-(8,6) nanotube complexes in macrophages treated with U18666A or Lalistat. **(b)** Overlay of transmitted light with hyperspectral image of RAW 264.7 macrophages incubated with ss(GT)_6_-(8,6) complex under the specified treatments. Color legend maps to nanotube emission peak wavelength. Scale bar = 50 µm. **(c)** Histogram of emission wavelengths of all pixels from the hyperspectral images, bin size of 1 nm. **(d)** Schematic of the optical setup for high-throughput measurements of ss(GT)_6_-(8,6) emission in live cells. **(e)** Spectra of ss(GT)_6_-(8,6) emission from live RAW 264.7 cells incubated in normal media (control), and in media with U18666A. **(f)** Mean center wavelengths of ss(GT)_6_-(8,6) emission from n = 5 technical replicates, acquired using hyperspectral imaging (I) and spectroscopy (S). Mean values were compared using a one way ANOVA with Sidak’s multiple comparison test. Error bars are standard error of the mean.

Using near-IR hyperspectral microscopy, we acquired the emission from the complexes localized within the endolysosomal organelles of macrophages under these three conditions (Figure 5a). When the emission wavelength is mapped to a color-scale and overlaid on a transmitted light image, the spatially-resolved emission from ss(GT)_6_-(8,6) complexes resulted in live-cell maps of endolysosomal lipid content (Figure 5b). The mean emission blue-shift for the two drug-treated conditions was over 11 nm compared with the control, with a single population for all three conditions, indicating lipid accumulation in all endolysosomal organelles that contained the complexes (Figure 5c). Neither Lalistat 3a2 nor U18666A directly perturbed the emission wavelength of the complexes (Figure S26). The endolysosomal lipid maps thus reflect the optical response of the complexes to the accumulation of lipids within the endolysosomal lumen. We quantified the nanotube emission response as a function of loading concentration. At a 5-fold dilution of the working concentration, the response of the emission wavelength to U18666A-induced lipid accumulation did not change, indicating that the function of the ss(GT)_6_-(8,6) complex was stable over a range of concentrations (Figure S27).

Based on the findings from our experiments, we present a model for the optical response of ss(GT)_6_-(8,6) ssDNA-nanotube complexes in live cells under the conditions investigated. The complexes enter the endolysosomal pathway and localize within the lumen of endolysosomal organelles. In the complex environment of the endolysosomal lumen, the nanotube emission functions as a cell-specific spectral fingerprint for the lipid content. Upon dysfunction of the NPC1 protein via introduction of U18666A to cells, free cholesterol (FC) accumulates within the lumen of the LE/Ly and adsorbs to the surface of the ss(GT)_6_-(8,6) complex, resulting in a distinct blue-shifting response of the nanotube emission. Similarly, upon introduction of Lalistat to cells, cholesteryl esters (CE) accumulate in the endolysosomal lumen, resulting in the adsorption of CE to the nanotube surface, and a concomitant blue-shifting response. In both situations, multiple secondary lipids also accumulate in the lysosomal lumen^53^. Hence, we find that the ss(GT)_6_-(8,6) DNA-nanotube complex functions as a quantitative optical reporter of lipid accumulation in the endolysosomal lumen via solvatochromic shifting of intrinsic carbon nanotube photoluminescence. Henceforth, we refer to ss(GT)_6_-(8,6) as the ‘reporter’.

As the nanotube emission is exclusively from the endolysosomal lumen, we investigated whether the nanotube spectra alone, obtained without imaging, could function as an analytical tool to benchmark drug-induced endolysosomal lipid accumulation in a high-throughput format. Using a customized nIR spectrometer^54^, we obtained spectra from complexes within live RAW 264.7 macrophages (Figure 5d) and detected a 9 nm shift in the emission from cells treated with U18666A –an example of a cationic amphiphilic drug which induces DIPL^55^ (Figure 5e). These results were similar to the hyperspectral data of control and U18666A-treated macrophages (Figure 5f), indicating the amenability of the complexes for both imaging and a spectroscopy-based assay which facilitates higher throughput.

### Measurement of lysosomal storage disorder phenotype in NPC1 patient-derived fibroblasts

The reporter was assessed for its response in primary cells from a patient with Niemann-Pick type C1 (NPC1), a lysosomal storage disease characterized by an accumulation of unesterified cholesterol in the lysosomes^56-57^. Fibroblasts collected from the NPC patient, as well as wild-type (WT) human fibroblasts, were incubated with the reporter for 30 minutes, then rinsed, and incubated in fresh media. Hyperspectral data collected after 24 hours indicates that the reporter blue-shifted by an average of 6 nm in NPC1 human fibroblasts, compared to WT human fibroblasts (Figure 6a). The lipid content of WT fibroblast endolysosomal organelles was relatively homogeneous, as evinced by the narrow distribution of the reporter emission (Figure. 6b). In contrast, the reporter exhibited a broad emission profile within NPC1 cells (Figure 6b), confirming the published findings that NPC1 cells exhibit a wide distribution of lipid concentrations^58^. The histogram also suggests that a sizeable fraction of endolysosomal organelles contain near-normal cholesterol content.

**Figure 6.**
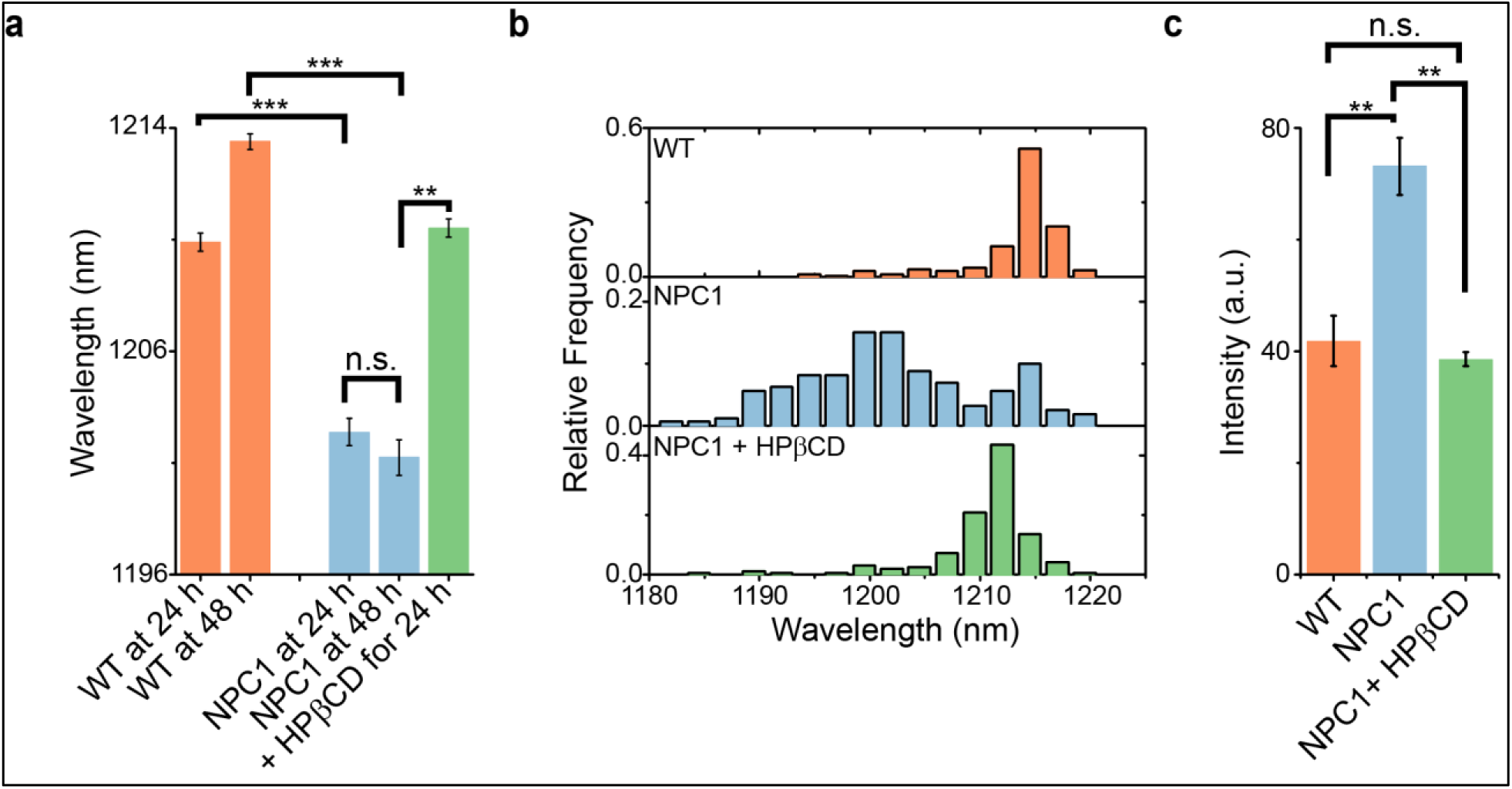
Measurement of endolysosomal lipid accumulation and reversal in NPC1 patient-derived fibroblasts. (a) Mean reporter emission from wild-type fibroblasts, patient-derived NPC1 fibroblasts, and NPC1 fibroblasts treated with hydroxypropyl-β-cyclodextrin (HPβCD) for 24 hours. Statistical significance was determined with a one-way ANOVA with Sidak’s multiple comparison test. **(b)** Histograms of the nanotube emission wavelength from single endolysosomal organelles of wild-type fibroblasts, NPC1 patient fibroblasts, and NPC1 fibroblasts treated with HPβCD for 24 hours. **(c)** Mean filipin intensity from WT fibroblasts, NPC1 cells, and NPC1 cells treated with HPβCD for 24 hours, at 48 hours after nanotube addition. Statistical significance was determined with a one-way ANOVA with Tukey’s multiple comparison test. All error bars are S.E.M. from 3 technical replicate experiments. * = p<0.05, ** = p<0.01, *** = p<0.001.

To test the reversibility of the reporter in live cells, we measured the emission from the reporter in NPC1 fibroblasts treated with hydroxypropyl-β-cyclodextrin (HPβCD), a drug known to facilitate the efflux of accumulated cholesterol from the lumen of endolysosomal organelles^59^. After treatment with 100 µM HPβCD treatment for 24 hours, hyperspectral imaging revealed significant red-shifting of the reporter emission from the same NPC1 cells that were previously blue-shifted (Figure 6a-b). The distribution of reporter emission from the HPβCD-treated NPC1 fibroblasts appeared notably similar to the emission from WT fibroblasts (Figure 6b), and reduction of total cell cholesterol content was confirmed by fixing the cells and staining with filipin (Figure 6c). Filipin staining was more pronounced in NPC1 fibroblasts as compared to WT fibroblasts, and HPβCD treatment resulted in significantly diminished filipin signal in NPC1 cells (Figure 6c, S28), thus validating the results obtained from the reporter alone. Additionally, pre-treatment of NPC1 fibroblasts with HPβCD prevented the blue-shifting of the reporter (Figure S29).

### Single-cell kinetic measurements of lipid accumulation

Recent technological advances have lead to an appreciation of the complexity of cell populations and the heterogeneity of single cells within a population of cells that seems uniform on a macroscopic level^60^. The ability of the developed reporter to function in live cell imaging applications uniquely positions it to provide data on a single cell level. Such data would have the potential identify previously unknown sub-populations of cells that could be crucial towards increasing our understanding of cellular lipid processing.

We assessed whether the reporter could measure single-cell kinetics of endolysosomal lipid accumulation. RAW 264.7 macrophages were cultured in media with lipoprotein depleted serum (LPDS).Under these conditions the reporter emission wavelength was approximately 1200 nm. Next, both AcLDL and U18666A were added to the cell media to induce endolysosomal lipid accumulation. Endolysosomal lipid accumulation was monitored via the acquisition of hyperspectral images every 10 minutes for 2 hours (Figure 7a). Single-cell emission trajectories were computed for all cells with reporter peak emission intensities of over 4 times higher than the background. The mean reporter emission from individual cells blue-shifted and reached equilibrium over approximately 90 minutes (Figure 7b). For each cell, the spectral trajectories were fit with a single exponential function to obtain the time constant, time lag, and the starting and final reporter wavelengths (Figure S30). The time constants of lipid accumulation in the cells averaged approximately 40 minutes and followed a lognormal distribution (Figure 7c). One potential explanation for this finding is that a lognormal distribution is often observed for a quantity arising from multiple serial processes^61^. We believe that this distribution of time constants from individual cells is consistent with the process of endolysosomal lipid accumulation, which involves sequential processes including LDL uptake into the early endosome, delivery of LDL to the lysosome, hydrolysis of esterified cholesterol by LAL, and the inhibited efflux of free cholesterol by NPC1.

**Figure 7.**
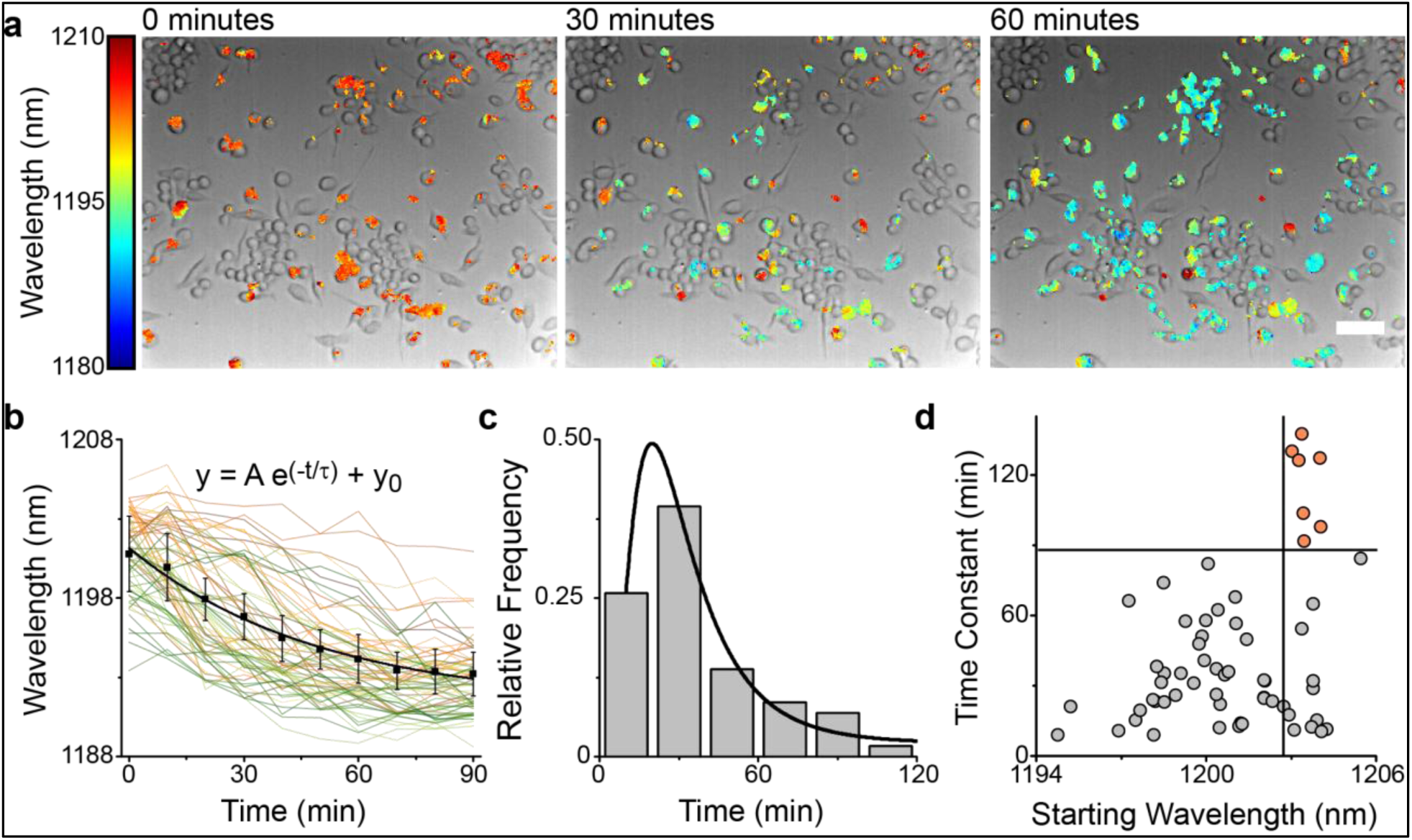
Single-cell kinetics of lipid accumulation. (a) Overlaid brightfield and hyperspectral images of the reporter emission in RAW 264.7 macrophages upon addition of AcLDL (100 µg/mL) and U18666A (3 µg/mL) to LPDS media at t = 0, 30 minutes, and 60 minutes. Scale bar = 50 µm. **(b)** Single-cell trajectories of lysosomal lipid accumulation from 60 cells. Black curve is the mean. Error bars are standard deviation from n = 4 technical replicates. **(c)** Distribution of time constants of lipid accumulation in single cells fitted to a log-normal distribution. Bin size = 20 nm. **(d)** Scatter plot of the time constants vs. starting emission wavelength, n = 60 cells.

Interestingly, the distribution of time constants showed marked heterogeneity, with the slowest and fastest time constants differing by an order of magnitude. Cells exhibiting slower rates of lipid accumulation also exhibited an initial state of relative lipid deficiency. The time constant of lipid accumulation modestly correlated with the starting wavelength (Spearman correlation of 0.33, p<0.01). Notably, the slowest shifting cells (defined as cells with a time constant > 90 minutes) exhibited an initial average emission wavelength >1202 nm, 2 nm more red-shifted than the faster shifting cells (Figure 7d). This result suggests the existence of a subpopulation of cells that maintained both lipid-deficient endolysosomal organelles and an extremely slow rate of lipid accumulation; i.e. these individual cells in the population may be especially resistant to endolysosomal lipid accumulation.

### Quantifying intracellular heterogeneity in monocyte endolysosomal organelles

After demonstrating the reporter’s ability to quantify lipid accumulation on a single cell level, we investigated whether the reporter could measure lipids on single-organelle level. We generated hyperspectral maps to quantify the intracellular heterogeneity of lipid content within individual endolysosomal organelles of single bone marrow derived monocytes cells during the differentiation process into bone marrow derived macrophages (BMDM) (Figure 8a). Hyperspectral images of two cells (Figure 8b) from the bone marrow-derived monocyte population on day 3 of differentiation showed distinctly different intracellular heterogeneity of the reporter response, evident from the histogram widths (Figure 8c). To mathematically quantify the intracellular heterogeneity of reporter emission within each cell, we used the normalized Simpson’s Index (nSI), a statistical measure of diversity^62-63^. For the cells shown in Figure 8c, the nSIs were significantly different (Cell 1 nSI = 0.18, Cell 2 nSI = 0.74). By plotting the mean reporter emission wavelength and nSI for a large population of cells on day 3, cells with the most lipid-rich endolysosomal organelles (shorter wavelength) exhibited greater intracellular heterogeneity in endolysosomal lipid contents (Figure 8d, Spearman correlation of 0.33, p < 0.01, for n = 64 cells). This observation, confirmed in BMDMs isolated from 4 different mice (Figure S31), suggests that within differentiating monocytes, cells with lipid-rich endolysosomal organelles maintain a sub-population of these organelles that are relatively lipid-deficient. Moreover, observation of BMDMs throughout differentiation validated the ability of the reporter to detect endolysosomal lipid accumulation in a non-pathological condition (Figure S32-33).

**Figure 8.**
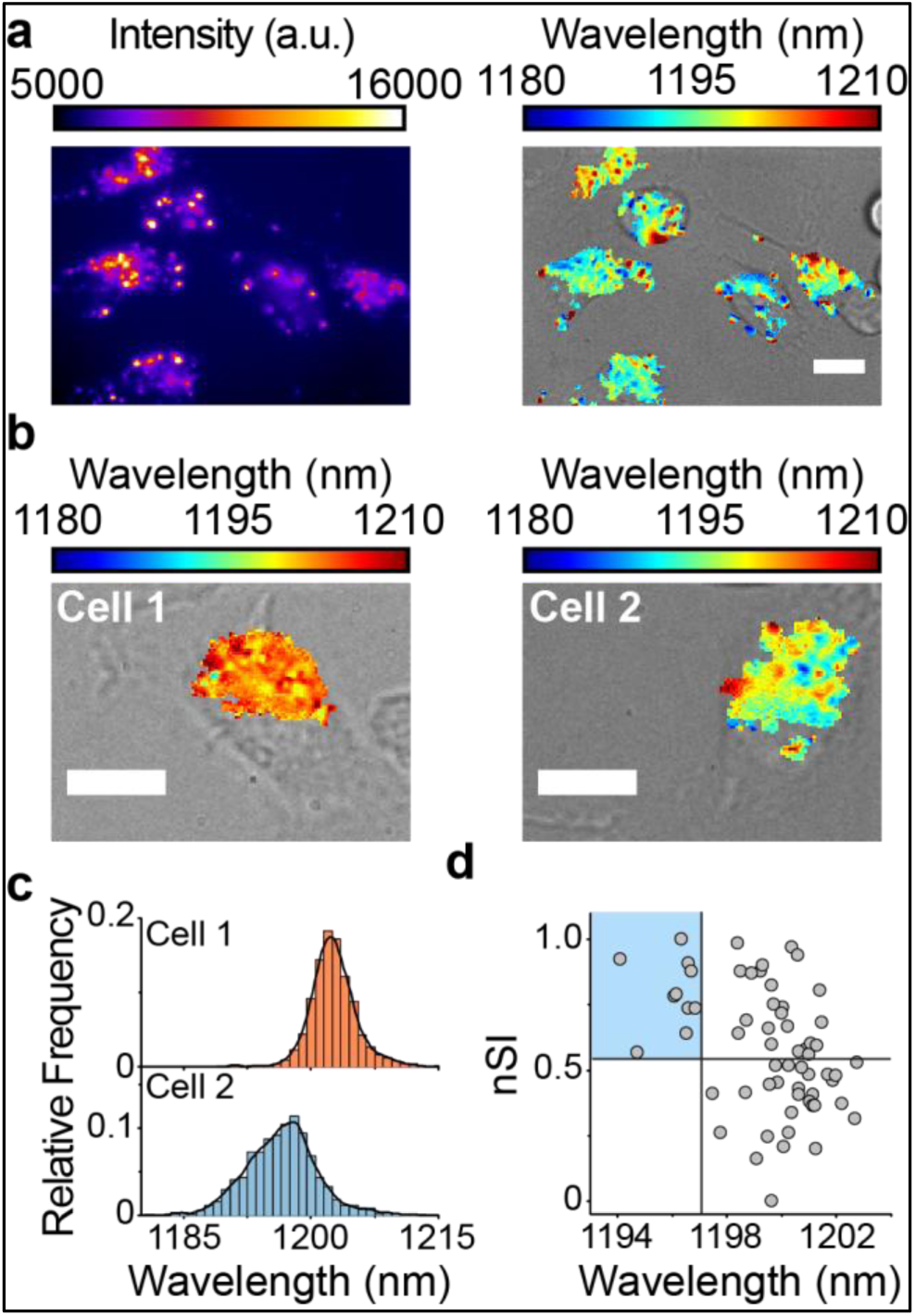
Quantifying intracellular heterogeneity in monocyte endolysosomal organelles. (a) Near-infrared broadband and overlaid brightfield/hyperspectral images of the reporter in bone marrow-derived cells after 3 days of colony stimulating factor-1 (CSF-1) treatment. **(b)** Endolysosomal lipid maps of the reporter in bone marrow-derived cells at day 3 of maturation. **(c)** Corresponding histogram of reporter emission from emissive pixels within the two cells. Bin size = 1 nm. **(d)** Scatterplot of the normalized Simpson’s Index (nSI) against the mean emission wavelength per cell for n = 64 cells. Scale bars = 10 µm

## Conclusion

In this work, we developed a carbon nanotube optical reporter, composed of the (8,6) SWCNT species non-covalently functionalized with a short DNA oligonucleotide, that can function as an optical reporter of lipid content within the endolysosomal lumen of live cells. The near-infrared photoluminescence of the nanotube responds quantitatively to lipids in the local environment of the reporter via shifting of the nanotube optical bandgap. Experimental evidence and all-atom molecular dynamics simulations suggest that the mechanism of the response is solvatochromic shifting due to the reduction in water density on the nanotube surface. The reporter remains within the endocytic pathway and localizes to the lumen of endolysosomal organelles without adversely affecting organelle morphology, structural integrity, or function. In the endolysosomal lumen, the reporter’s near-infrared emission responds rapidly and reversibly to lipid accumulation. Via nIR hyperspectral microscopy, the reporter can quantitatively map lipids in live cells. Using spectroscopy, the reporter can measure endolysosomal lipid accumulation in live cells in a high-throughput drug screening-type format. The reporter detected lipid accumulation in lysosomal storage disorders, including in the endolysosomal organelles of fibroblasts derived from a patient with Niemann-Pick Type C, as well as phenotypic reversal in the same cells after drug treatment. The reporter functioned with single-cell and single-organelle resolution and was used to assess single cell kinetics of modified LDL accumulation within endolysosomal organelles, showing that the rate of cholesterol accumulation differs by an order of magnitude across macrophages in the same population of cells. Endolysosomal lipid accumulation in differentiating bone marrow-derived monocytes was also observed, and high resolution endolysosomal lipid maps revealed intracellular heterogeneity in the form of a sub-population of lipid-deficient endolysosomal organelles in lipid-rich cells. As the first technique for measuring lipid flux in the endolysosomal lumen of live cells, we expect this tool will have broad utility in both drug screening applications and in the investigation of disease pathways associated with altered lipid biology such as atherosclerosis, neurodegenerative diseases, lysosomal storage disorders, and liver disease.

## Materials and Methods

### DNA encapsulation of single-walled carbon nanotubes

The chemical reagents were purchased from Sigma-Aldrich (St. Louis, MO, US) and Fisher Scientific (Pittsburgh, US). Single-walled carbon nanotubes (SWCNTs) produced by the HiPco process were used throughout the study (Unidym, Sunnyvale, CA). The carbon nanotubes were dispersed with DNA oligonucleotides via probe-tip ultrasonication (Sonics & Materials, Inc.) of 2 mg of the specified oligonucleotide (IDT DNA, Coralville, IA) with 1 mg of raw SWCNT in 1 mL of 0.1 M NaCl for 30 minutes at 40% of the maximum amplitude of the ultrasonicator (SONICS Vibra Cell). Following ultrasonication, the dispersions were ultracentrifuged (Sorvall Discovery 90SE) for 30 minutes at 280,000 x *g*. The top 3/4 of the resultant supernatant was collected and its concentration was determined with a UV/Vis/nIR spectrophotometer (Jasco, Tokyo, Japan) using the extinction coefficient Abs_910_ = 0.02554 L mg^-1^cm^-1^ ^36^. To remove free DNA, 100 kDa Amicon centrifuge filters (Millipore) were used to concentrate and re-suspend the DNA-nanotube complexes.

### Purification of single-chirality nanotube complexes

Carbon nanotubes were separated by an ion-exchange chromatography (IEX) method according to the procedure described by Tu et al^32^. Briefly, unsorted HiPco nanotubes were dispersed using a DNA oligonucleotide with the sequence ss(GT)_6_, as described above. The sample was injected into a high performance liquid chromatograph (HPLC) (Agilent, 1260 Infinity) fitted with an anion-exchange column (Biochrom Labs, Inc., CNT-NS1500) with a running buffer of 2x SSC at a flow rate of 2 mL/min. A linearly increasing salt concentration gradient of 1M NaSCN (5%/min) was used to elute the nanotubes from the stationary phase and fractions were collected. The first fraction exciting the HPLC contained the highest purity of the (8,6) species, estimated at 86%, which was used for subsequent studies.

### Near-infrared fluorescence microscopy of single-walled carbon nanotubes

As described in a previous study^36^, near infrared fluorescence microscopy was used to acquire the photoluminescence emission from SWCNTs. The system comprised a continuous wave (CW) 730 nm diode laser with an output power of 2 W injected into a multimode fiber to produce the excitation source for fluorescence experiments. To ensure a homogenous illumination over the entire microscope field of view, the excitation beam passed through a custom beam-shaping module to produce a top-hat intensity profile with under 20% power variation on the imaged region of the sample. The final power at the sample was 230 mW. A longpass dichroic mirror with a cut-on wavelength of 875 nm (Semrock) was aligned to reflect the laser to the sample stage of an Olympus IX-71 inverted microscope (with internal optics modified to improve near-infrared transmission from 900-1400 nm) equipped with a 20X LCPlan N, 20X/0.45 IR objective and a UAPON100XOTIRF, 1.49 oil objective (Olympus, USA). Emission was collected with a 2D InGaAs array detector (Photon Etc.). Z-stacks of broadband, widefield emission (from 900 to 1650 nm) were acquired in steps of 0.5 µm, which satisfied the Nyquist sampling criterion for 100X magnification. Custom codes, written using Matlab software, were used to subtract background, correct for non-uniformities in excitation profile, and compensate for dead pixels on the detector. Hyperspectral microscopy was conducted by passing the emission through a volume Bragg grating (VBG) placed immediately before the InGaAs array in the optical path. The filtered image produced on the InGaAs camera was composed of a series of vertical lines, each with a specific wavelength. The reconstruction of a spatially-rectified image stack was performed using cubic interpolation on every pixel for each monochromatic image, according to the wavelength calibration parameters. The rectification produced a hyperspectral “cube” of images of the same spatial region exhibiting distinct spectral regions with 3.7 nm FHWM bandwidths. Approximately 50% of the emission intensity from the nanotubes is reduced after passage through the VBG.

### Analysis and processing of hyperspectral data

Hyperspectral data acquired was saved as a (320 x 256 x 26) 16-bit array, where the first two coordinates signify the spatial location of a pixel and the last coordinate is its position in wavelength space. For the (8,6) nanotube, the 26-frame wavelength space ranges from 1150 nm to 1250 nm. An initial filter removed any pixels with a maximum intensity value outside (1170 to 1220 nm), as these were background pixels that emit outside the range for (8,6). For the remaining pixels, a peak-finding algorithm was used to calculate the intensity range for a given pixel i.e. range = (intensity_maximum – intensity_minimum). A data point was designated as a peak if its intensity was range/4 greater than the intensity of adjacent pixels. Pixels that failed the peak-finding threshold, primarily due to low intensity above the background, were removed from the data sets. The remaining pixels were fit with a Lorentzian function. Fits with R^2^ < 0.8 were also removed.

### Preparation of nanotubes labeled with visible fluorophores

To increase the fluorescence intensity of the Cy3 or Cy5 fluorophores attached to DNA strands encapsulating SWCNTs, a 6 nucleotide long polyT tail was added to the end of the (GT)_6_ sequence, as fluorophores near the surface of SWCNTs are known to quench^64^ (Integrated DNA Technologies, sequence = GTGTGTGTGTGTTTTTTT). For confocal imaging with Alexa-647 SWCNT, a small polyethylene glycol spacer was also added to further increase the fluorescence intensity of the fluorophore (Integrated DNA Technologies, sequence = GTGTGTGTGTGTTTTTTT/iSP18//3Alexaf647N//3’). These modified DNA-strands were non-covalently complexed with HiPco SWCNTs via the previously described sonication and centrifugation protocol.

### Preparation of gold nanoparticle-conjugated nanotubes

Gold nanoparticle-conjugated nanotubes were prepared according to a previously published study. Briefly, 10 nm citrate-capped gold nanoparticles were synthesized by bringing 50 mL of 0.01 wt% HAuCl4 and adding 2 mL of 1 wt% Sodium(III) citrate. After 2 minutes, the solution turns bright red, indicating nanoparticle formation. The gold nanoparticles were stabilized via a ligand exchange reaction by shaking overnight with bis(p-sulfonatophenyl)phenylphosphine dihydrate dipotassium salt. The nanoparticles were then centrifuged and resuspended in deionized water. In parallel, ss(GT)27-T6-thiol-dispersed HiPco nanotube complexes were created by means of the previously described sonication and centrifugation protocol. The nanotube complexes were filtered with ultracentrifuge filters three times to remove unbound DNA. The nanotube complexes and excess gold nanoparticles were then shaken overnight. The unbound gold nanoparticles were removed via centrifugation, which would pellet the unbound nanoparticles, and careful supernatant extraction.

### Transmission electron microscopy (TEM) imaging

Gold nanoparticle-nanotube conjugates were first imaged on carbon coated TEM grids by letting a 20 µL drop evaporate in the center of the grid. For imaging in RAW 264.7 cells, gold nanoparticle-nanotubes were introduced to the media for 30 minutes at 1 mg/L, and then washed thoroughly and replaced with fresh media. After 6 hours, cells were washed with serum-free media then fixed with a modified Karmovsky’s fix of 2.5% glutaraldehyde, 4% paraformaldehye and 0.02% picric acid in 0.1M sodium cacodylate buffer at pH 7.2. Following a secondary fixation in 1% osmium tetroxide and 1.5% potassium ferricyanide, samples were dehydrated through a graded ethanol series, and embedded in an epon analog resin. Ultrathin sections were cut using a Diatome diamond knife (Diatome, USA, Hatfield, PA) on a Leica Ultracut S ultramicrotome (Leica, Vienna, Austria). Sections were collected on copper grids and further contrasted with lead citrate and viewed on a JEM 1400 electron microscope (JEOL, USA, Inc., Peabody, MA) operated at 120 kV. Images were recorded with a Veleta 2K x2K digital camera (Olympus-SIS, Germany).

### Fluorescence microscopy of live cells

Standard fluorescence imaging in the UV-visible emission range was performed on the hyperspectral microscope by using an XCite Series 120Q lamp as the light source and a QiClick CCD camera (QImaging) directly attached to a c-mount on a separate port of the microscope. Fluorescence filter sets from Chroma Technology and Semrock were used. Confocal imaging was performed on a Zeiss LSM 880, AxioObserver microscope equipped with a Plan-Apochromat 63× Oil 1.4 NA differential interference contrast (DIC) M27 objective in a humidified chamber at 37 °C. Z-stacks were obtained using a step size of 198-220 nm.

### Fluorescence spectroscopy of carbon nanotubes in solution

Fluorescence emission spectra from aqueous solutions of SWCNTs were acquired using a home-built apparatus consisting of a tunable white light laser source, inverted microscope, and InGaAs nIR detector^54^. The SuperK EXTREME supercontinuum white light laser source (NKT Photonics) was used with a VARIA variable bandpass filter accessory capable of tuning the output 500 – 825 nm with a bandwidth of 20 nm. During the course of the measurements, the excitation wavelength remained at 730 nm, close to the resonant excitation maximum of the DNA-encapsulated (8,6) nanotube species. The light path was shaped and fed into the back of an inverted IX-71 microscope (Olympus) where it passed through a 20x nIR objective (Olympus) and illuminated a 100 µL nanotube sample at a concentration of 0.2 mg/L in a 96-well plate (Corning). With an exposure time of 1 second, the emission from the nanotube sample was collected through the 20x objective and passed through a dichroic mirror (875 nm cutoff, Semrock). The light was f/# matched to the spectrometer using several lenses and injected into an Isoplane spectrograph (Princeton Instruments) with a slit width of 410 µm which dispersed the emission using a 86 g/mm grating with 950 nm blaze wavelength. The spectral range was 930 – 1369 nm with a resolution of ∼0.7 nm. The light was collected by a PIoNIR InGaAs 640 x 512 pixel array (Princeton Instruments). A HL-3-CAL-EXT halogen calibration light source (Ocean Optics) was used to correct for wavelength-dependent features in the emission intensity arising from the spectrometer, detector, and other optics. A Hg/Ne pencil style calibration lamp (Newport) was used to calibrate the spectrometer wavelength. Background subtraction was conducted using a well in a 96-well plate filled with DI H_2_O. Following acquisition, the data was processed with custom code written in Matlab which applied the aforementioned spectral corrections, background subtraction, and was used to fit the data with Lorentzian functions.

### Nanotube chirality and DNA sequence-dependent response to LDL

Unsorted DNA-SWCNT samples were diluted to 2 mg/L in PBS and incubated with 0.5 mg/mL LDL (Alfa Aesar) for 18 hours at 37 °C. Chirality-separated samples were diluted to 0.2 mg/L in PBS and incubated with 0.5 mg/mL LDL for 18 hours at 37 °C. Controls were incubated with no LDL present. Photoluminescence spectra were acquired with 2-second exposure times.

### Titrations of DNA-nanotube complexes with PEG-conjugated lipids

Unsorted ss(GT)_6_-DNA-SWCNT samples were diluted to 2 mg/L in PBS and incubated with various concentrations (0-5µM) of two polyethylene glycol (PEG)-conjugated lipids (Cholesterol-PEG 600, “Cholesterol-PEG”, Sigma Aldrich; C16 PEG750 Ceramide, Avanti Lipids). Samples were incubated for 2 hours at 37 °C. Photoluminescence spectra were acquired with 2-second exposure times under 730 nm laser excitation. Polyethylene glycol, with molecular weights of 600 or 750 kDa, diluted in PBS, were used as controls to test for non-specific interactions. To test the effect of lowered pH on sensor performance, samples were diluted in a 100 mM pH 5.5 acetate buffer instead of PBS.

### Calculation of normalized Simpson’s Index

The Simpson’s index is a diversity index used to measure the richness and evenness of a basic data type^62^. The diversity index is maximized when all types of data are equally abundant. When applied to microbiology, the Simpson’s Index is referred to as the Hunter-Gaston index^63^. In our application,

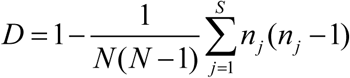

where D is the diversity index, N is the total number of pixels within each cell with detectable nanotube emission, S is the total number of histogram bins, and n_j_ is the total number of pixels within the j^th^ bin. We obtained the Simpson’s Index for each cell, SI_j_. For the set of SIj calculated for all the cells in an experiment, we normalized the value obtain the normalized Simpson’s Index, nSI.

### Cell culture reagents and conditions

RAW 264.7 TIB-71 cells (ATCC, Manassas, VA) were grown under standard incubation conditions at 37°C and 5% CO_2_ in sterile, filtered DMEM with 10% heat-inactivated FBS, 2.5% HEPES, 1% glutamine, and 1% penicillin/streptomycin (all from Gibco). For studies performed with homozygous mutant NPC, compound mutant heterozygote NPC, or wild-type fibroblasts, the respective cell lines, GM18453, GM03123 or GM05659 (Coriell, Camden, NJ) were cultured in MEM with 10% FBS, 2.5% HEPES, and 1% glutamine. Cells were plated on glass bottom petri dishes or lysine-covered glass dishes (MatTek) for fibroblasts. U2OS-SRA cells Chirality-separated SWCNTs were added at 0.2 mg/L in cell culture media (70 µL total volume) and incubated with cells for 30 minutes at 37 °C. This corresponds to approximately 0.5 picogram of SWCNT per cell, for a 50% cell confluency in a 9 mm diameter glass-bottom dish. Cells were imaged immediately, or trypsinized (Gibco), and re-plated on a fresh petri dishes before hyperspectral imaging. All cells were used at 50-70% confluence.

### Filipin staining of NPC1 patient-derived fibroblasts

Cells were fixed with 4% paraformaldehyde for 15 minutes, washed 3x with PBS, and stained with Filipin III (Sigma) at a concentration of 50 mg/L for 20 minutes. The cells were then washed 3x with PBS and imaged using a DAPI filter cube.

### Cell viability and proliferation assays

RAW 264.7 macrophages were seeded in untreated 96-well plates at 7,000 cells per well. The reporter was introduced to the cells at 0.2 mg/L. Reporter, vehicle (0.027 MSSC + 0.1 M NaSCN), or hydrogen peroxide-treated cells were incubated (times indicated), washed, and detached from the plate with Versene (1x PBS without Mg^2^+/Ca^2+^, 5mM EDTA, 2% FBS), pelleted, and incubated with Annexin V Alexa Fluor and Propidium Iodide (Life Technologies). Cells were analyzed by imaging cytometry (Tali) to quantify cell number and fluorophore content. For proliferation assays, RAW 264.7 macrophages treated with 0.2 mg/L ss(GT)_6_-(8,6)-SWCNT or vehicle were seeded at 150,000 cells on 100 mm diameter untreated culture dish on day zero. After settling for 10 hours, cells were harvested with Versene (1x PBS without Mg^2+^/Ca^2+^, 5mM EDTA, 2% FBS) and by mechanical tapping to remove cells from the surface, stained with Calcein AM, and counted for the initial seeding density. Media was replaced every 2 days. At each 24 hour period, cells were harvested as before and counted. Cell counts represent Calcein AM positive (live) cells.

### Lipidomics analyses

Six T-175 flasks were seeded at <50% confluence with RAW 264.7 macrophages in their fifth passage. Three flasks each were treated as follows: control (untreated) or carbon nanotube treated (0.2 mg/mL for 30 minutes, washing, and incubating for 6 hours). Cells were harvested (approx. 80% confluence in each set of flasks) by manual scraping. The flask contents corresponding to each condition were pooled into separate conical tubes and spun to pellet, washed in PBS containing protease inhibitor cocktail (Thermo-Pierce, 88666), pelleted, and flash frozen in a dry ice/IPA slurry with a small head volume of the PBS/inhibitor.

For cell fractionation we adapted a sucrose/iodixanol equilibrium gradient centrifugation procedure for lysosome separation (Thermo, 89839). Briefly, flash frozen cells were thawed and suspended in 2x cpv of PBS/inhibitor, vortexed with reagent included in the kit, and Dounce homogenized with a cooled, tight fitting pestle using 90 strokes (starting cell material was >500mg). After homogenization, reagent was added, tube inverted, and then centrifuged at 4°C, 500x rcf for 10 minutes. The pellet was stored as the debris/nuclear fraction in all experiments. The supernatant was taken for subsequent ultracentrifugation. Briefly, one day before ultracentrifugation, a sucrose/iodixanol gradient (bottom to top: 30, 27, 23, 20, 17%) was layered into 12 mL polyallomer tubes (Thermo, 03699) and allowed to equilibrate in a cold room inside the metal buckets of an appropriate hanging-bucket rotor. The supernatant from above was mixed with the sucrose/iodixanol gradient to make a final sample density of 15% (total volume, 1 mL), which was then gently layered onto the top of the pre-formed gradient. The buckets were then sealed and moved into the ultracentrifuge using the following settings: ∼135,000 rcf (32,000 rpm), 2.5 hours running time, acceleration/deceleration 9/5, 4°C. The fractionated cell supernatant was removed from the top, and pipetted into six fractions based on volume removed from the ultracentrifuge tube. The volume removed from top to bottom was kept constant across the three conditions. Fractions were frozen until analysis.

Each of the six fractions, plus the nuclear/debris fraction (7 total/condition) was analyzed for the three conditions (21 fractions total). To quantify total protein, a standard curve was produced using BSA mixed into the sucrose/iodixanol gradient (Bradford assay background versus varying gradient was not different). 1x Bradford reagent at room temperature was mixed 1:1 with standard, allowed to incubate in the dark for 30 minutes, and the absorption measured at 595 nm. Each of the fractions was analyzed in this manner after addition of 0.2% IPEGAL CA-630 (non-ionic detergent) to solubilize bound proteins.

Cholesterol quantification was performed (Sigma, MAK043) using a coupled enzyme reaction between cholesterol oxidase and peroxidase with a proprietary colorimetric probe. Cholesterol esterase was used before the reactions to ensure total cholesterol was measured. Briefly, a 3-phase extraction was performed on each sample fraction (7:11:0.1 chloroform:isopropanol:IPEGAL CA-630). The top aqueous phase and interphase were removed, and the bottom organic phase was vacuum dried. The dried fractions were resuspended in provided buffer and reacted for 1 hour at 37°C with the supplied reagents and the absorption read at 570 nm. This was compared to a standard curve. Cholesterol levels were normalized by total protein content as measured by the Bradford assay.

Total lipid (total unsaturated hydrocarbon, including cholesterol) was measured after chloroform extracting the samples as above. Briefly, a phospho-vanillin color producing reagent was made by dissolving 5 mg/mL vanillin (Sigma, V1104) in 200 µL neat ethanol, and adding this to the appropriate volume of 17% phosphoric acid. This reagent was stored cool in the dark until needed. Dried sample fractions in glass vials were processed as follows: to each vial was added 200 µL ∼98% sulfuric acid. The dried contents were coated with the acid by tipping the vial and vortexing. The vial was placed into a 100°C mineral oil bath for 20 minutes. The resulting brown/black material was rapidly cooled in wet ice slurry for at least 5 minutes, and 100 µL was placed side-by-side into a 96-well glass plate. To one well, 50 µL of the phospho-vanillin reagent was added, mixed, and the plate incubated in the dark for 12 minutes. Absorption of each well (reagent reacted and sulfuric acid background) was taken at 535 nm. The difference was the measurement, which was compared to a standard curve that used Oleic acid (Sigma, O1008), prepared using the above protocol, as a model unsaturated hydrocarbon material. Total lipid levels were normalized by total protein content.

### Extraction and differentiation of bone marrow-derived macrophages

Bone marrow derived cells (BMDM) were prepared from 6 week-old C57/Bl6 mice and cultured in the presence of 10 ng/ml of recombinant colony stimulating factor-1 (CSF-1)^65^. Cells were collected 3, 5 and 7 days post-isolation and submitted to flow cytometry analysis for expression of the differentiation markers Gr-1 (monocytes/granulocytes-1/200), Cd11b (macrophages-1/200), F4/80 (mature macrophages-1/50). Cells were incubated with 1 µl of Fc Block (BD Biosciences) for every million cells for at least 15 minutes at 4°C. Cells were then stained with the appropriate antibodies (BD Biosciences) for 20 minutes at 4°C, washed with FACS buffer, and resuspended in FACS buffer containing DAPI (5 mg/ml diluted 1:5,000) for live/dead cell exclusion^66^.

### LysoTracker-nanotube colocalization

RAW 264.7 or BMDM macrophages were incubated with Cy5-ss(GT)_6_-HiPco nanotubes for 30 minutes at a concentration 1 mg/L. The cells were then washed 3x with PBS and placed in fresh cell media. Six hours later, the cells were incubated with 5 nM LysoTracker Green DND-26 (Life Technologies) for 15 minutes in cell media, washed 3x with PBS, and imaged immediately in fresh PBS. The FITC or Cy5 channels were used for LysoTracker Green or Cy5-ss(GT)_6_-HiPco nanotubes, respectively. Plates of cells containing only Cy5-ss(GT)_6_-HiPco nanotubes or LysoTracker Green were used as controls to test for bleed-through across channels.

### Atomic force microscopy (AFM)

A stock solution of ss(GT)_6_-(8,6)-SWCNTs at 7 mg/L in 100 mM NaCl was diluted 20x in dH_2_O and plated on a freshly cleaved mica substrate (SPI) for 4 minutes before washing with 10 mL of dH_2_O and blowing dry with argon gas. An Olympus AC240TS AFM probe (Asylum Research) in an Asylum Research MFP-3D-Bio instrument was used to image in AC mode. Data was captured at 2.93 nm/pixel XY resolution and 15.63 pm Z resolution.

### Statistics

Statistical analysis was performed with GraphPad Prism version 6.02. All data met the assumptions of the statistical tests performed (i.e. normality, equal variances, etc.). Experimental variance was found to be similar between groups using the F-test and Brown-Forsythe test for unpaired t-tests and one way ANOVAs, respectively. To account for the testing of multiple hypotheses, one way ANOVAs were performed with Dunnet’s, Tukey’s, or Sidak’s post tests when appropriate. Sample size decisions were based on the instrumental signal-to-noise ratios.

### Cell line source and authentication

RAW 264.7 and U2OS cells were acquired from ATCC, and were tested for mycoplasma contamination by the source. Primary bone-marrow derived monocytes were tested for mycoplasma contamination using Dapi staining. Patient-derived fibroblasts were obtained from Coriell and tested for mycoplasma contamination by the source.

### Code availability

Matlab code for the data analysis in this manuscript is available upon request, by contacting the corresponding author.

## ASSOCIATED CONTENT

### Supporting Information

The supporting information contains supplementary figures (S1-33), videos (1-3) and a supplementary table.

## Notes

The authors declare no competing financial interest.

## Acknowledgements

This work was supported by the NIH Director’s New Innovator Award (DP2-HD075698), NIH grants R37-DK27083, R01CA148967, R01CA181355, P20-GM103430 and P30 CA008748 cancer center support grant, the Rhode Island Foundation (20164347) the Anna Fuller Fund, the Louis V. Gerstner Jr. Young Investigator’s Fund, The Expect Miracles Foundation - Financial Services Against Cancer, the Experimental Therapeutics Center, Mr. William H. Goodwin, Mrs. Alice Goodwin, the Commonwealth Foundation for Cancer Research, the Honorable Tina Brozman Foundation, the Alan and Sandra Gerry Metastasis Research Initiative, Cycle for Survival, the Frank A. Howard Scholars Program, and the Center for Molecular Imaging and Nanotechnology at Memorial Sloan Kettering Cancer Center. P.V.J. was supported by an NIH NCI-T32 fellowship (2T32CA062948-21). D.R. was supported by an American Cancer Society 2013 Roaring Fork Valley Research Fellowship. C.P.H. was supported in part by National Cancer Institute (NCI) Grant NIH T32 CA062948, J.M. was supported by the U.S. Department of Energy (DOE), Office of Science, Basic Energy Sciences (BES) under Award DE-SC0013979. L.A. was supported by the American Brain Tumor Association. J.B. was supported by a Tow Fellowship Award from the Center for Molecular Imaging and Nanotechnology, Memorial Sloan Kettering Cancer Center. We thank the Molecular Cytology Core Facility at Memorial Sloan Kettering Cancer Center and the Electron Microscopy & Histology Core Facility at Weill Cornell Medical College. Use of the high-performance computing capabilities of the Extreme Science and Engineering Discovery Environment (XSEDE), supported by the National Science Foundation (NSF) grant numbers TG-MCB-120014 and TG-MCB-130013, is gratefully acknowledged. We also thank Y. Shamay, A. Erez, R. Williams, R. Langenbacher, and J. Shah for helpful discussions, and J. Bartlett for aid in manuscript preparation.

---

FOR TOC ONLY

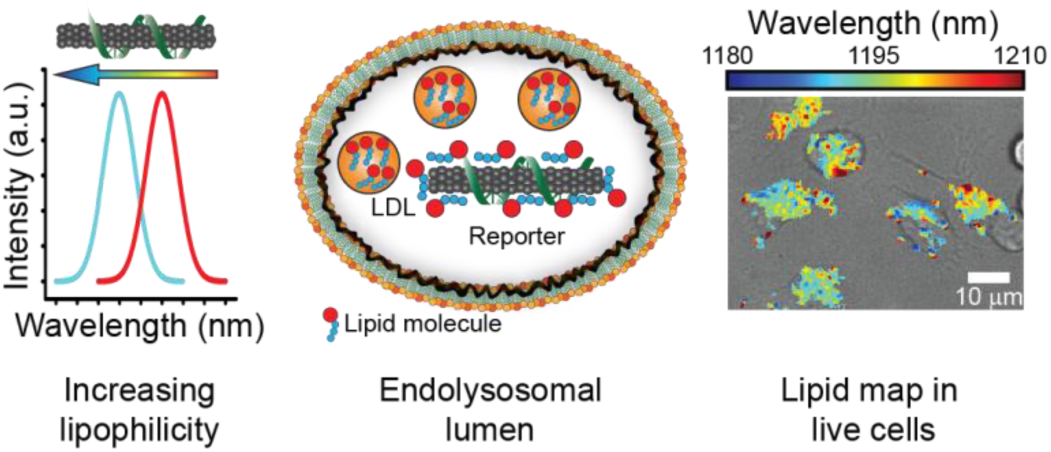

